# Structural basis of transcription: RNA Polymerase II substrate binding and metal coordination at 3.0 Å using a free-electron laser

**DOI:** 10.1101/2023.09.22.559052

**Authors:** Guowu Lin, Christopher O. Barnes, Simon Weiss, Bercem Dutagaci, Chenxi Qiu, Michael Feig, Jihnu Song, Artem Lyubimov, Aina E. Cohen, Craig D. Kaplan, Guillermo Calero

## Abstract

Catalysis and translocation of multi-subunit DNA-directed RNA polymerases underlie all cellular mRNA synthesis. RNA polymerase II (Pol II) synthesizes eukaryotic pre-mRNAs from a DNA template strand buried in its active site. Structural details of catalysis at near atomic resolution and precise arrangement of key active site components have been elusive. Here we present the free electron laser (FEL) structure of a matched ATP-bound Pol II, revealing the full active site interaction network at the highest resolution to date, including the trigger loop (TL) in the closed conformation, bonafide occupancy of both site A and B Mg^2+^, and a putative third (site C) Mg^2+^ analogous to that described for some DNA polymerases but not observed previously for cellular RNA polymerases. Molecular dynamics (MD) simulations of the structure indicate that the third Mg^2+^ is coordinated and stabilized at its observed position. TL residues provide half of the substrate binding pocket while multiple TL/bridge helix (BH) interactions induce conformational changes that could propel translocation upon substrate hydrolysis. Consistent with TL/BH communication, a FEL structure and MD simulations of the hyperactive Rpb1 T834P bridge helix mutant reveals rearrangement of some active site interactions supporting potential plasticity in active site function and long-distance effects on both the width of the central channel and TL conformation, likely underlying its increased elongation rate at the expense of fidelity.

## Introduction

An array of structural studies has revealed critical details of the mechanisms of multisubunit RNA polymerases (msRNAPs)^1-4^, of which RNA Polymerase II (Pol II) is the eukaryotic enzyme responsible for protein coding gene transcription. msRNAP activity functions in an iterative nucleotide addition cycle of substrate selection, catalysis, and movement of one base (translocation)^5,6^. Evidence supports that this nucleotide addition cycle is undertaken in part by rearrangements of critical, highly-conserved structural domains termed the “bridge helix” and “trigger loop”^7-14^. From the very first structural models of msRNAPs, attention was drawn immediately to the bridge helix. The factors for this included the conservation of bridge helix sequence, its prominent and striking positioning adjacent to the nucleotide addition site, and its presence as an obstacle around which template bases must pass to enter the substrate-binding site. This latter observation, coupled with distinct BH conformations observed in eukaryotic and prokaryotic structures led to suggestions of bridge helix motions participating in translocation steps^12-14^.

Matching of an incoming nucleotide (NTP) to its corresponding Watson-Crick pair on the template strand is first step in DNA-mediated transcription by Pol II. NTP-matching triggers or captures conformational changes in the highly conserved Rpb1 trigger loop (TL) residues (1077-1096) leading to NTP stabilization, phosphodiester bond formation, and pyrophosphate release^11^. X-ray diffraction structures have shown the endpoints of these states, namely TL residues away from (“off” conformation) or forming part (“on” conformation) of the NTP binding pocket along with potential intermediates^9,11,15-17^. Biochemical, biophysical, and genetic studies support TL control of substrate specificity and catalysis^10,18-27^. Computational studies suggest that the TL functions in positioning the incoming NTP^28,29^ and have provided suggestions on how dynamics of the TL and BH may participate the elongation cycle^30-32^. It was originally proposed that the TL might function in acid-base catalysis through an ultra-conserved histidine (Rpb1 His^1085^ in yeast)^11^, in a mechanism potentially conserved among nucleic acid polymerases^33^. Subsequent studies illustrated that catalysis could be supported by residues as diverse as glutamine or leucine in place of the conserved histidine, suggesting no need for TL residues in acid-base catalysis or that it could be bypassed^18,19,23,24^. The simplest interpretation is that the TL acts as a positional catalyst for matched substrates associated with correctly positioned Mg^2+^ ions. Capturing the exact catalytic conformation of TL with substrate has been difficult because complexes must be compromised either by use of non-hydrolyzable substrate analogues or chain-terminated 3′-deoxy RNA templates in order to prevent catalysis^11,15,34^. Furthermore, no closed/“on” TL conformations have been described for any msRNAP since the *T. thermophilus* substrate bound complex was described in 2007^9^.

In addition to the lack of high-resolution studies describing the Pol II active site during transcription, no msRNAP structures clearly accommodate the two active site Mg^2+^ ions proposed to act universally by Steitz and colleagues in nucleic acid polymerases^35^. The initial tour de force studies from the Kornberg lab observed that the second metal, “Metal B” was low occupancy and was not placed in the structure of apo-Pol II^36^. For Pol II elongation complexes especially, observation of two metals likely has been limited by lower resolution, the difficulty in capturing the TL in a closed/”on” state, and presumptive radiation damage (during X-ray data collection), which can trigger decarboxylation of aspartate and glutamate residues and hence compromise metal coordination. Understanding the arrangement of critical active site features is important in its own right, but also critical to interpret how mutations alter Pol II activity.

Experiments first in yeast and then in other organisms illustrated that Pol II activity can be modulated by mutation, either increasing or decreasing catalytic activity in vitro and in vivo. Alterations to catalytic properties can alter Pol II fidelity and overall elongation rate^20,25-27,37-42^, and in doing so alter Pol II dynamics on the template and co-transcriptional processes^43-49^. The molecular basis of Pol II mutant infidelity in at least one case has best been understood as relating to altered TL dynamics^26^, however how mutants might alter active site configuration leading to changes in activity are not understood at a structural level. We have previously identified a Pol II BH proline substitution (*rpb1* T834P) that confers hyperactivity^23^. Specific BH proline substitutions in both yeast and archaeal polymerases can confer hyperactivity and are interpreted as potentially favoring a kinked/altered BH conformation that might normally occur during msRNAP nucleotide addition cycles^7,8,23^. Additionally, other BH substitutions in *E. coli* RNAP also confer increased activity above WT, consistent with perturbations to BH function being able to increase catalytic activity^50^. We reasoned critical details of the Pol II active site orientation might be revealed by revisiting Pol II substrate addition by X-ray crystallography at higher resolution and by extending analysis to a the hyperactive T834P Pol II enzyme predicted to kink the BH.

During crystallography experiments at the synchrotron, the catalytic Mg of Pol II are susceptible to X-ray induced photo-reduction and disorder. To obtain an uncompromised structure of the Pol II active site metal centers, we performed experiments at the Linac Coherent Light Source (LCLS) X-ray Free Electron Laser (XFEL). The bright ultra-short X-ray pulses (∼40 fs) produced by the LCLS enable a ‘diffraction-before-destruction’ approach to record diffraction images before meaningful radiation damage has time to occur^51^. Because the process destroys the sample volume exposed, the sample must be continually replaced, a method termed Serial Femtosecond Crystallography (SFX). We performed fixed-target SFX experiments using a highly automated goniometer-based instrumentation^52,53^ to understand the process of nucleotide addition by Pol II for both the wild type enzyme and the Rpb1 T834P BH mutant.

Our studies show clear density confirming the occupancy of two catalytic Mg^2+^ ions, and evidence for a third putative catalytic Mg^2+^ within the Pol Il’s active site. We also observed structural details of the TL at the highest resolution yet achieved. In addition, the structure of the hyperactive bridge helix mutant T834P and molecular simulations suggest how this mutant compromises transcriptional fidelity, how BH and TL contacts can be altered by altered BH conformation, and potential pathways for the BH to influence both the TL and Pol II cleft width at a distance.

## Results

Two key technical advancements have allowed us to obtain WT and mutant Pol II structures that capture Pol II in a substrate bound, closed TL “on” conformation. First, we explored new crystallization conditions and identified a promising new, high salt condition allowing good diffraction (Methods). Second, we applied SFX methods at the LCLS XFEL source. Our earliest SFX experiments used a type of liquid/crystal injector, the gas-dynamic virtual nozzle (GDVN), to introduce a stream of Pol II nanocrystals (of between ∼0.1 - 5 µm) through the X-ray beam position^54^. However, because of the shear forces involved, Pol II crystals are not good candidates for GDVN delivery; crystal fragility limited diffraction results to only a small number of low resolution images, 4 Å resolution at best^55^. We next turned to a fixed-target approach for SFX utilizing a goniometer setup installed at the LCLS XPP instrument, and later incorporated into the LCLS Macromolecular Femtosecond Crystallography instrument (MFX)^52,56^]. For fully automated experiments, a SAM automounter^57^ robotically exchanges samples to/from a precision high-speed goniometer, which rapidly rotates and translates crystals into the beam position^52^. Using this setup, we employed a highly efficient and reliable data collection strategy, centered on the use of large area (1.5 mm x 1.5 mm) Mylar support grids. These grids can accommodate dozens of crystals and are compatible with UV fluorescence microscopy for crystal imaging and identification^53,58^. UV microscopy imaging was used to map the location and size of the individual crystals on grids and this information was fed into automated routines incorporated into the controls systems of the standard goniometer setup at MFX. Employing these routines, the goniometer rapidly positioned crystals, distributed in random locations on the grids, into the X-ray beam position, which resulted in highly accurate crystal targeting with indexing rates better than 70% [Barnes, 2019]. This procedure allowed collection of a 3.0 Å data set of WT Pol II from 220 crystals (**Supp Fig. 1A**). Diffraction data was processed using the cctbx.xfel software tools^59^ and the structure was solved by molecular replacement using WT Pol II as search model in Molrep^60^. Refinement was performed using Phenix ^59^ to a final R_cryst/_R_free_ (%) of 28.3/26.2 respectively (**Table 1**), the resultant electron density map was of high quality and allowed residue placement (**Fig. 1A**). At the initial stages of refinement, we observed electron density in the difference map (Fo-Fc) for the matched ATP, all residues of the TL (**Supp. Fig. 1B**), Mg^2+^ ions (sites A and B), and of great interest, the putative third catalytic Mg^2+^ “site C” (**Fig. 1B)**, observed previously (in a similar position) in DNA Pol η^61^, proposed to be critical for catalysis^62^. A third Mg^2+^ has also been observed for DNA Pol β, but was suggested to bind after chemistry^63^ (see Discussion).

**Figure 1.**
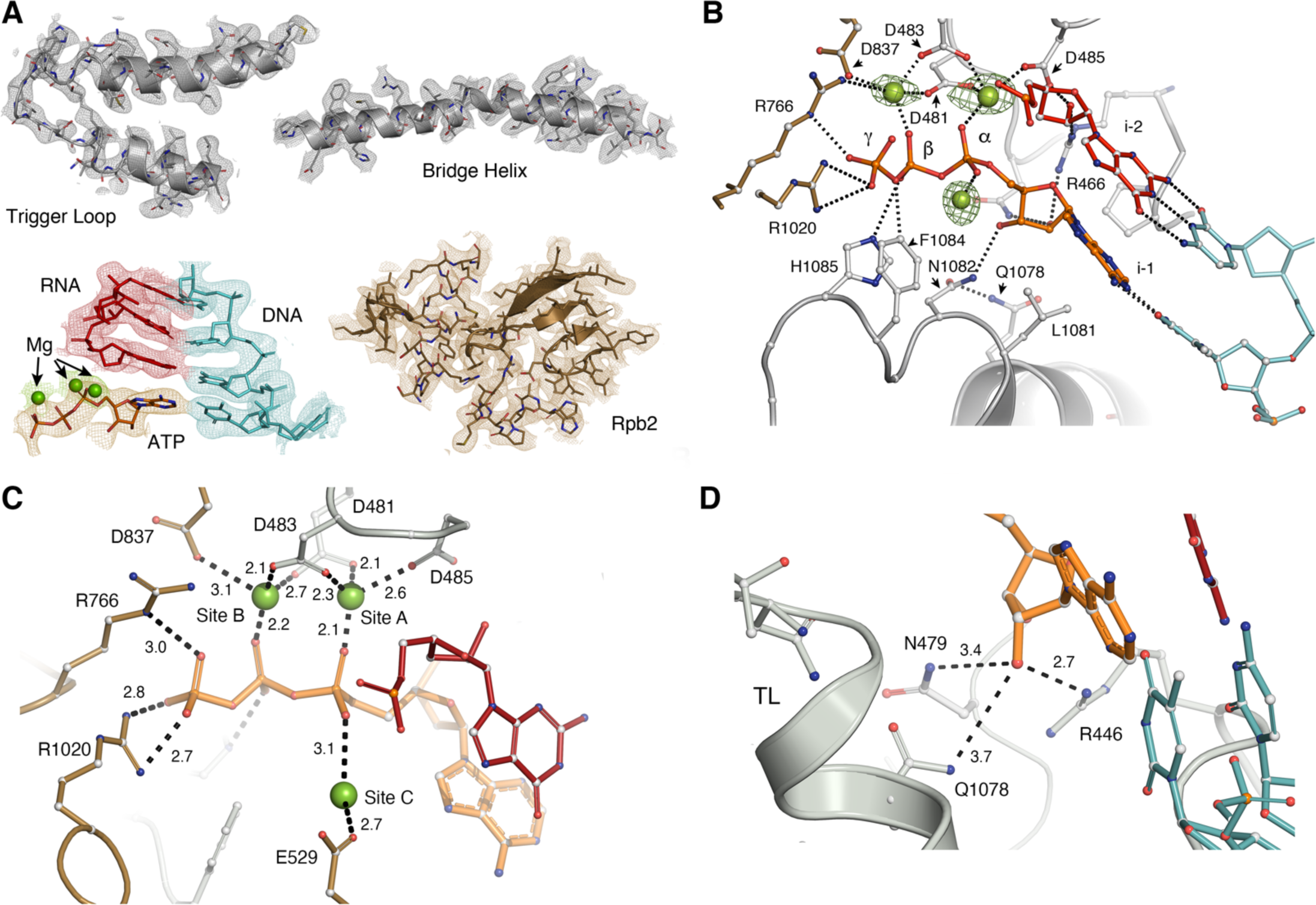
Features of Pol II XFEL structure at 3.0Å. **A.** The following coloring scheme will be used throughout the manuscript, Rpb1 residues, silver, Rpb2 residues, gold, template strand (cyan), nascent mRNA strand (red), ATP substrate, orange, magnesium ions, green. Electron density maps (2Fo-Fc) for Rpb1 trigger loop and bridge helix; Rpb2 residues 792-845 the nucleic acid scaffold, including the template strand the nascent mRNA strand, the ATP substrate, and the three Mg^2+^ ions. **B.** Cartoon and ball stick representation illustrating the mechanism of ATP stabilization in Pol II’s active site; green mesh, electron density (Fo-Fc, difference map) contoured at 6 σ around the three Mg^2+^ ions. **C.** Coordination of the three Mg^2+^ atoms by Rpb1 P-loop residues Asp^481,483^ ^and^ ^485^, and Rpb2 Asp^837^; coordination distances (in angstroms) are indicated next to dashed lines. **D.** Rpb1 residues involved in selection of substrate 2′-OH group.

### The radiation-damage free FEL structure of Poll II reveals the presence of three metals in the nucleotide binding pocket

Questions regarding positioning of the full metal content of Pol II’s active site in an elongation complex have been outstanding since no clear electron density for both site A and B could be resolved at the low resolution of the data sets^11^, as noted by^9^ in their supplemental discussion (referring to MgI (A) and MgII (B)). A putative Mg^2+^ B was observed in the apo-Pol II structure but not placed in the model due to low occupancy^36^. Our radiation damage-free XFEL data at 3.0 Å resolution illustrates unambiguously the presence of site A and site B metal ions (with similar occupancies and B-factors) and their coordination in Pol II’s active site (**Fig. 1B and C**). The inner coordination shell of site A and B metals comprises Rpb1 (P-loop) acidic residues Asp^481^, Asp^483^, and Asp^485^; Rpb2 Asp^837^; and the α and β phosphate oxygens from ATP (**Fig. 1C**). The outer coordination shell comprising Rpb1 Arg^446^, and Rpb2 Arg^1020^ and Lys^987^ stabilizes the inner shell and could play a role in enhanced metal affinity^64^ (**Supp Fig. 1C**). The modeled inter-magnesium distance is 3.9 Å, which is comparable to the bacterial RNA Polymerase (RNAP) (3.9 Å)^9^ and the DNA Pol η (3.5 Å)^61,62^ hence site A and B metal coordination patterns vary only slightly between the three enzymes, with the caveat that 3′-deoxy chain termination (**Supp. Fig. 1D**, this study) can distort active site geometries. To observe bimetallic coordination in the presence of a 3′-OH RNA we soaked Pol II crystals with ATP at several concentrations of manganese chloride (MnCl_2_) and collected X-ray data using conventional synchrotron radiation. Electron density for site A was observed at 5mM Mn^2+^, however, no electron density was observed for site B at this concentration. Density for both sites was observed at high concentrations of the metal, 250 mM MnCl_2_; however, no ATP binding was detected at this Mn^2+^ concentration (**Supp. Fig. 1E**). Thus, in the presence of an RNA 3′-OH group, the interatomic site A to site B distance is 3.8 Å. Occupancy refinement of site A was slightly higher than that of site B, consistent with the former being more fully coordinated (**Supp Fig. 1E).**

Of great interest in the XFEL structure was the presence of positive electron density for a putative third catalytic Mg^2+^. Refinement of a water molecule placed inside site C density yielded low B-factor values (18 Å^2^) while refinement with a Mg^2+^ atom had similar values (68 Å^2^) as those in metals A (52 Å^2^) and B (81 Å^2^). Thus, the observed density may correspond to a third, putatively catalytic Mg^2+^ ion (site C). Site C stabilization (**Fig. 1C**) is achieved through contacts with the α-phosphate (3.2 Å) and with Rpb2 E529 (3.1 Å). Moreover, molecular dynamics simulations show that site C metal is stable at this position and tends to oscillate around the α-phosphate, Rpb2 E529 and the phosphodiester bond of the i-2 base (4.2 Å away) (**Supp. Fig. 1F** and G). To corroborate that the use of XFEL was essential to observe the three metals in the active site, we collected conventional synchrotron X-ray data (beamline 12-2 SSRL) from the same crystal batch at 3.3 Å resolution; as expected, the difference map of the refined data, where all metals were omitted from the calculation showed electron density for site A only, however the TL was still visible in the closed “on” position supporting our new crystallization condition as an advance over previous conditions where the closed TL has been exceedingly difficult to observe (**Supp. Fig. 1H**).

### TL interactions in the active site

Our XFEL structure provides the first reproduction of a closed Pol II TL conformation (TL_C_) since 2006^11^ and does so in a robust fashion at a considerable increase in resolution. Importantly, the higher resolution and more advanced refinement programs allow a fresh look at the position of U-shaped TL_C_ relative to substrate and other Pol II active site residues (**Fig. 2A and B).**

**Figure 2.**
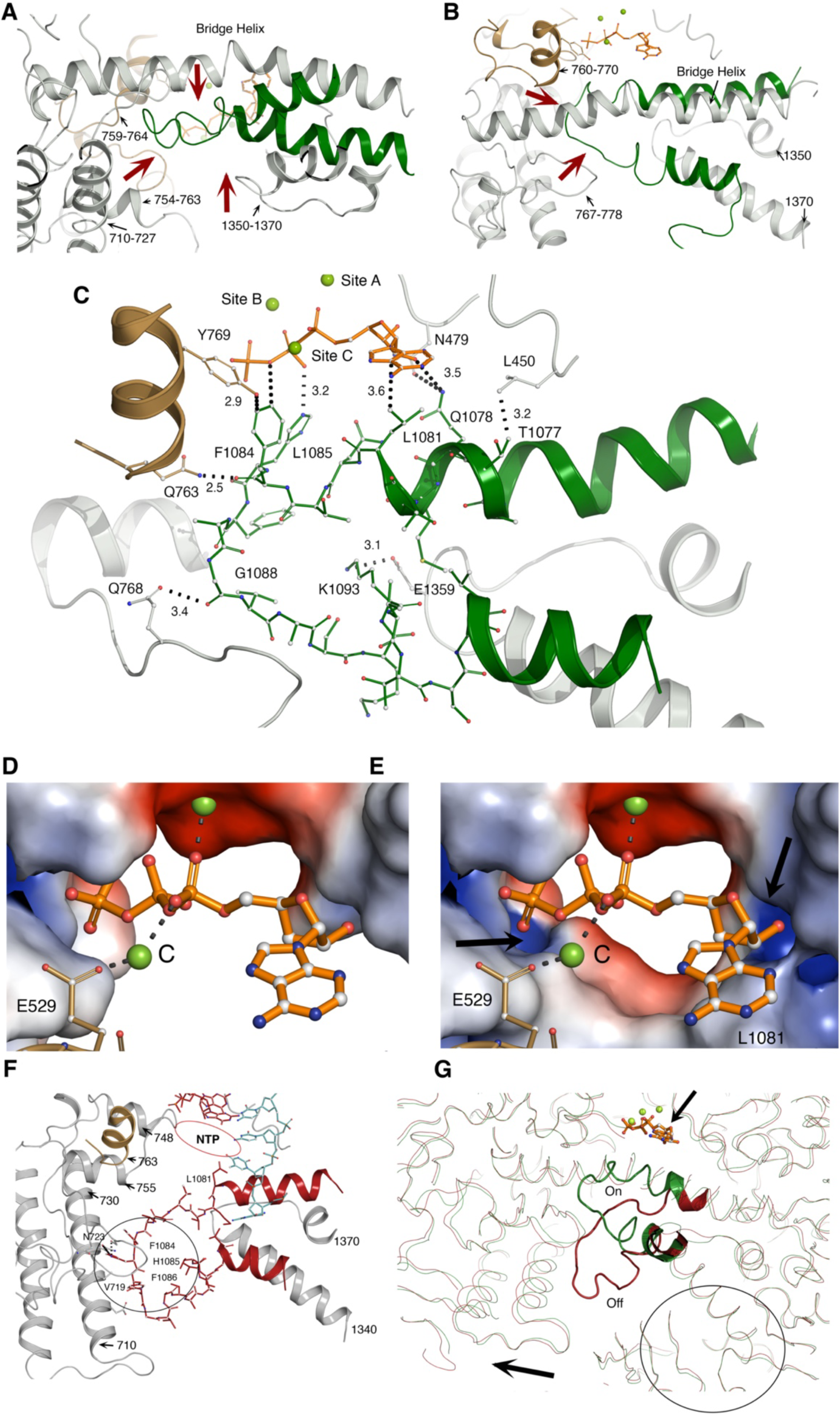
Formation of substrate binding pocket by Pol II Trigger Loop closure. **A and B** Cartoon and ball and stick representation of TL residue (illustrated in green) packing (in the closed state) by neighboring Rpb1 regions. **C.** Detailed view of key interactions that stabilize the close state. **D and E.** Surface electrostatic potential calculation using (using the APBS suite in Pymol) of Pol II’s active site in the open and closed TL conformations illustrating positive potential areas around the phosphate tail and the 2′-OH group (black arrows). Acidic (negative) surfaces shaded in red and basic (positive) surfaces in blue. **F.** Structure of Pol II’s active site without a NTP substrate illustrating the TL in the open conformation (black circle). **G.** Ribbon representation of the overlay between TL’s open (red) and closed (green) conformations illustrating the large-scale structural re-arrangements between the two states (black arrow). An ATP substrate is illustrated in orange (small arrow).

Four contact surfaces for the TL can be identified: the NTP side (residues 1078-1085), the tip (residues 1086-1089), the bridge helix face, and a small contact region (opposing the bridge helix face) with loop Rpb1 residues 1350-1365. The NTP side of the TL forms part of Pol II’s active site and contributes to ATP binding (**Fig. 1B and 2C**): Leu^1081^ forms stacking interactions with the purine ring as observed previously^11^. Gln^1078^ and Asn^1082^ form H-bonds with the 3′-OH of the ribose ring, and Phe^1084^ and His^1085^ interact with the phosphate tail (C-H and H-bonds with the β phosphate). Calculation of the electrostatic potential of the ligand-bound active site with the TL in the closed conformation and without it, shows that the TL residues form half of the NTP binding pocket that together with Arg^446^ and Asn^479^ form a positive complementary surface for the 2′-OH ribose moiety (critical for NTP vs dNTP selectivity) and the phosphate tail (**Fig 1D, 2D and E**). Asn^479^ has been previously implicated in 3′-OH recognition^34,65^, while Arg^446^ is positioned to interact with either 2′- or 3′-OH groups. It has been recently suggested that the bacterial residue homologous to Arg^446^ interacts productively with 2′-OH of the incoming NTP ribose, but non-productively with the 3′-OH of 2′-dNTPs, promoting selection of NTPs over dNTPs^66^. Here we observed for the first time Pol II Arg^446^ positioned to select the 2′-OH. Comparisons of the TL position and its surroundings in the opened (**Fig. 2F**) (TL_O_) and closed (TL_C_) conformations reveal a defined physical path that appears to guide its motion during the elongation cycle (**Supp Fig. 2B**). This physical path comprises Rpb1 residues 711-724 and 766-773 for the open form (**Fig. 2F**); and 819-838 (BH), Rpb2 763-766 and 1018-1021 for the closed form (**Figs. 2C and 4B**). The importance of these interactions for both states is manifested by gain/loss of function mutations (G/L-OF) which putatively alter TL mobility and thus the elongation cycle^23,25-27,43,67-70^. Moreover, it can be observed that residues neighboring the TL undergo significant “breathing motions” during the TL cycle suggesting dynamics of these regions may influence TL motion or states and may function as potential conduits of distal allosteric inputs into the TL (**Fig. 2G**).

**Figure 3.**
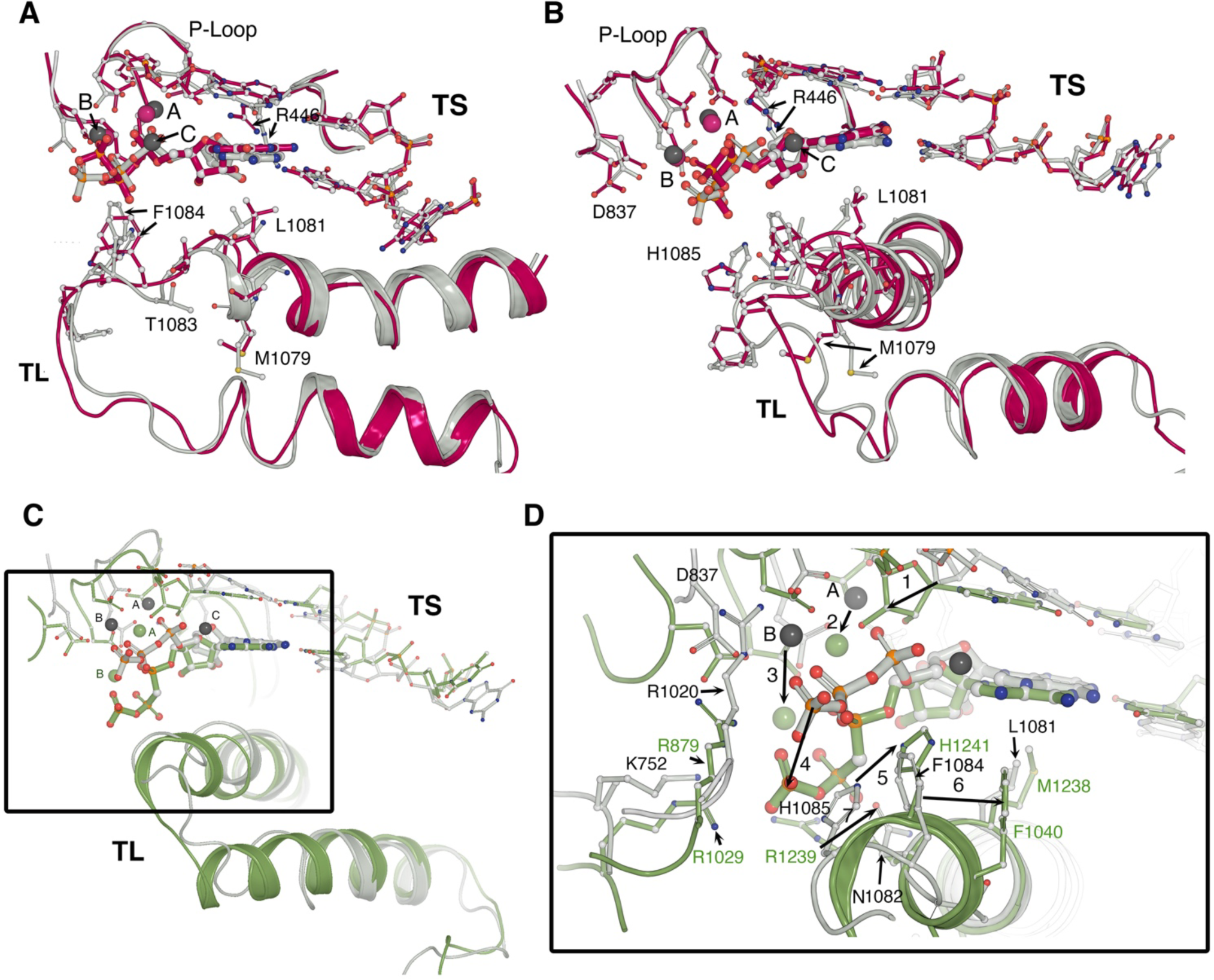
Comparison of closed Pol II Trigger Loop conformation determined by XFEL with previously published closed TL conformations. **A-B.** Two views of the overlay between WT XFEL structure (silver) and our refinement of PDB:ID 2E2H structure. Technological advances in recently developed refinement programs allowed higher confidence in positions and density for putative Mg^2+^ ions and side chain orientations in the XFEL- and 2E2H-determined TL conformations. **C-D.** Differences between eukaryotic Pol II substrate-bound (closed) TL conformation and *Tth* RNAP substrate-bound closed conformation complex.

**Figure 4.**
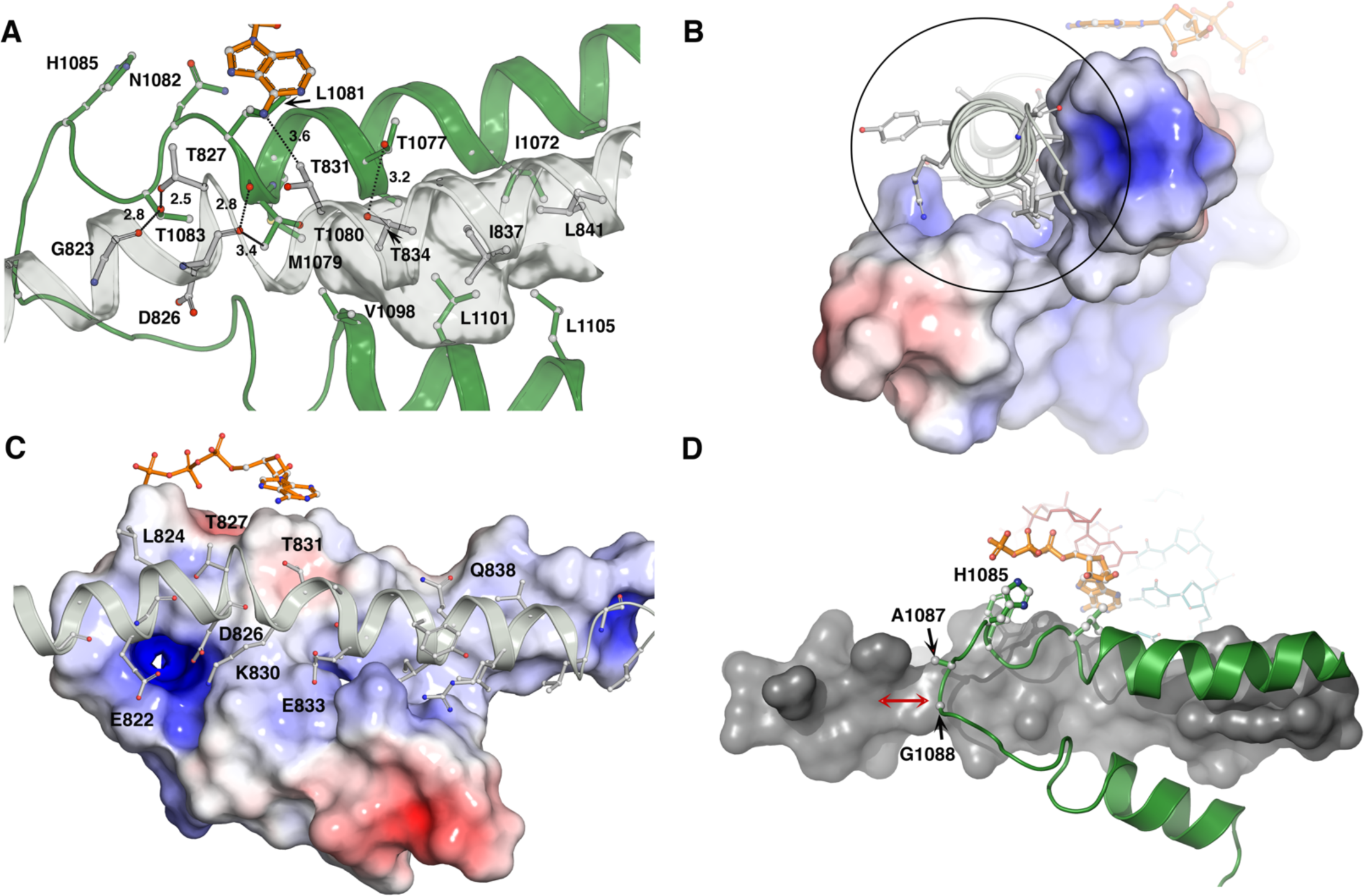
Network of interactions between the Pol II Trigger Loop and Bridge Helix. **A.** In the closed state the TL and BH residues form an interaction network, including H-bonds and a small hydrophobic pocket to accommodate BH residues Thr^834^, Ile^837^ and Leu^841^. **B-C**. Two views of TL-bridge helix interactions. Electrostatic surface representation of TL residues (calculated using the APBS suite in Pymol, ref) contacting bridge helix residues (shown as ball and sticks) illustrating charged BH-TL interactions. **D.** Short non-bulky residues, Ala^1087^ and Glly^1088^ (cartoon and ball and stick representations) at the tip of the TL are required to allow full motion of the TL over BH residues (shown as surface representation).

Comparisons of our new structure with the single closed TL structure described for Pol II (*S. cerevisiae* Pol II PDB 2E2H^11^)(**Fig, 3A-B)** and the single closed TL structure described for RNAP (*T. thermophilus* PDB 2O5J^9^)(**Fig. 3D-E)** show a number of differences. The presence of Mg^2+^ ions and their clear coordination was noted above, but comparison of structures also indicates differences with previously proposed positions. Different positioning of residue side chains can now be modeled with higher confidence due to increased resolution and technological advances in refinement approaches. A previously noted difference between open and closed TL conformations was the apparent release of Met^1079^ from the hydrophobic pocket/interface that enables packing of TL N- and C-helices with Rpb1 α46^16^. A number of increased- or putatively increased-activity substitutions^23,24,69,71^ affect this pocket led to a proposal that Met^1079^ release might be a key event in transition between open and closed states influenced by NTP binding^16^. In contrast, here we observe that the closed TL state is in fact compatible with retention of Met^1079^ within the hydrophobic pocket. Re-refinement of 2E2H suggests that side chain positions in our higher resolution data are also consistent with the electron density maps of 2E2H (**Supp Fig 3A and B**).

Comparison of the Pol II closed TL with the *Tth* (PDB:id:2O5J)^9^ closed TL structure shows similar geometries (**Supp Fig. 3B**), however several features are of note (**Fig. 3C-D**). The Pol II structure is more compact with shorter NTP and Mg distances suggesting the possibility that the *Tth* non-hydrolyzable analog may be in more of a pre-insertion state. Conversely, the non-hydrolyzable substrate and/or the RNA 3′-OH in the *Tth* structure and the chain-terminated state of the Pol II structure may allow different compaction. Interactions observed between TL and substrate in each structure are similar, though the triphosphates of the *Tth* structure are positioned differently in the less compact state. The *Tth* TL has Arg^1239^ which locates under the beta and gamma phosphates forming up to three H-bonds with the phosphate tail. This is in contrast to Pol II Asn^1082^ (at the analogous position) which locates 5 Å away from the tail; thus, these side chains do not make equivalent contacts with the substrate, consistent with potential plasticity in residue functions over evolution. Overall, the number of positively charged residues contacting the substrate is larger in the *Tth* structure creating an enhanced substrate affinity.

### Bridge helix residues lock the TL in the closed conformation while TL residues position substrate in a catalytic conformation

Interactions between the closed TL and the BH are extensive, including electrostatic and hydrophobic interactions. Bridge helix Gly^823^, Asp^826^, Thr^827^, and Thr^834^ form hydrogen bonds with TL residues (**Fig. 4A**). Moreover, TL_C_ residues form a shallow hydrophobic pocket to accommodate BH residues Ile^837^ and Leu^841^ (**Fig. 4A**). Calculation of TL_C_ surface electrostatic potential shows the presence of charged regions that foster interactions with BH charged residues (**Fig. 4B and C**). Mutagenesis studies of the TL shown that replacing Ala^1087^ and Gly^1088^ –which locate at the tip of the TL within 4.4 Å of BH residues (**Fig 4D**)– by bulkier residues^23^ are lethal in *S. cerevisiae*, underscoring the importance of a defined TL_C_ conformation to facilitate NTP positioning. A catalytic role for the TL has been proposed previously^11,72^, however, genetic and biochemical studies in both yeast and *E. coli* have indicated that mutations of His^1085^ (the putative catalytic residue, acting as a general acid) for bulky polar (glutamine) or even hydrophobic (leucine) can support growth and catalysis^18,19,23^.

### Structural Basis for enhanced activity of the bridge helix mutant T834P

Genetic and biochemical studies by our group and others have indicated higher elongation rates for mutants that could promote conformational changes in the BH^7,23^. One of these mutants, T834P has been shown to produce a ∼3-fold increase in yeast Pol II elongation rate in vitro and is hypersensitive to excess Mn^2+^ presence in the growth medium, which is a characteristic of fast elongating Pol II mutants that are defective for fidelity^23,73^. To understand the structural basis for such enhanced processivity we solved the FEL structure of the T834P mutant to 3.1 Å resolution. At this resolution, density for the mutated proline residue was distinguished unambiguously. Overlay of the mutant versus WT structures show slight BH rearrangements (R.M.S between BH residues 820-844 is 0.47) however conformational changes can be detected up to 40 Å away as result of disruptions of hydrogen bond networks that confer alternative, but putatively still enzymatically active conformations. Such changes include: 1) disruption of the H-bond (3.0 Å) between Thr^834^ and TL^1077^ (allosterically) drives Arg^446^ away from its position (critical for 2′-OH substrate discrimination) to form a H-bond with Asp^485^ (**Fig. 5A, arrows**). 2) Slight changes in metal A and B positions as well as NTP binding are observed (**Fig. 5B**), however, more significant are changes in metal C coordination. As consequence of a one angstrom change in the position of the BH around Ala^828^, the loop harboring Rpb2 Glu^529^, which coordinates Metal C (**Fig. 1C and Fig 5B**), shifts away from the coordination sphere of site C. On the other hand, Metal C in the mutant structure locates closer to the α-phosphate (2.2 Å). Biochemical studies of Rpb2 Glu^529^ ^70^ have shown that alanine or aspartate mutations (which would disrupt an interaction with metal C) are associated with faster elongation. Interestingly, substitution of Rpb2 Tyr^769^ to phenylalanine also results in faster elongation^68^. While it remains to be determined if these substitutions function through altered positioning of Metal C, it is intriguing that the two residues closest to Metal C are both implicated in shaping Pol II catalysis. 3) since the small hydrophobic pocket formed by the TL (to accommodate BH residues in the closed state) shrinks; and the number of H-bonds in the TL-BH interaction in the mutant is smaller, it is possible that the flexibility of the TL in the mutant is larger than WT (**Fig. 5C**). The latter has been associated biochemically and computationally with enhanced catalytic activity^74^. 4) A positively charged patch formed by Asn^479^, Gln^1078^ and Arg^446^ that interacts with the ribose 2′-OH in WT residues is disrupted in the mutant (**Fig. 5D**), thus possibly contributing to the enhanced dNTP misincorporation (**Supp. Fig 4B**). Finally, 5) as noted above, changes in structure are propagated far from the site of the substitution, including a widening of the Pol II cleft and altered conformation to TL residues adjacent to the C-terminal helix (**Fig. 5E**). The widening of the cleft may both enhance TL flexibility or reduce stabilization of the open TL conformation contributing to both hyperactivity and loss in fidelity of the mutant. MD simulations of the mutant support the enhanced TL flexibility, relevance increased distances and more potential openness in the static structure of Pol II T834P (**Supp. Fig 4C-E**). We also note that MD simulations indicate that Metal C is stably associated with this structure and appears to be coordinated by Rpb2 Tyr^769^ (**Supp. Fig. 4F**). Subtle but consistent increases in distance between the N-terminal portion (nucleotide interacting) of the TL and the BH are observed in T834P relative to WT while greater distances are observed between the BH and a distal TL residue (V1094). The T834P TL also exhibits altered flexibility (wider ranges of distances), including in the TL tip (1088-1099) relative to the BH.

**Figure 5.**
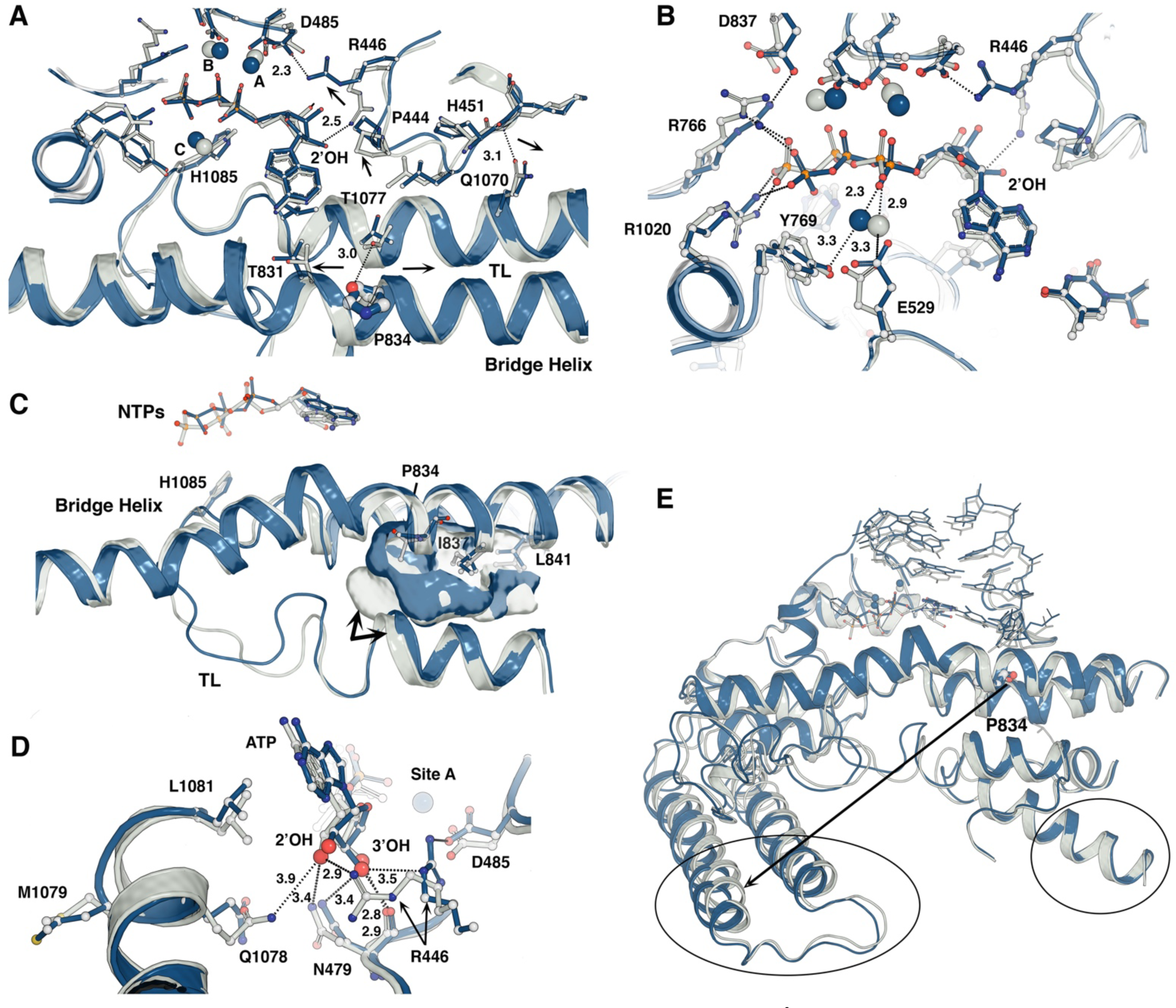
XFEL structure of hyperactive Pol II T834P mutant at 3.1Å reveals alterations at multiple scales. Silver WT, Blue T834P. A. Disruption of Thr^834^-Thr^1077^ H-bond triggers active site, BH and TL conformational changes. Repositioning of Arg^446^ away from the ribose 2′-OH results in loss of NTP vs dNTP discrimination. **B.** Overlay between WT and T834P active sites illustrating changes in active site geometry that impact metal C coordination. **C**. Overlay between WT and T834P TL structures show geometry changes that affect TL-BH interactions including a shallower hydrophobic pocket (see Fig. 4A) that could contribute to enhanced TL mobility. **D.** WT active site residues including Asn^479^ and R^446^ form H-bonds with ribose 2′-OH atoms, such interactions are lost in the T834P mutant resulting in decreased NTP-dNTP discrimination. **E.** Overlay between WT and T834P structures show that the T834P mutation triggers long-range allosteric structural changes (arrow and ellipse).

## Discussion

Structural studies of Pol II have revealed tremendous amounts of information about the organization of the Pol II complex, including the ever-increasing complexity of assemblages that function at individual steps in transcription as well as linking Pol II to cotranscriptional RNA processing events and the transit of Pol II through chromatin. At the fine scale, precise arrangement and dynamics of active site structures underlying chemistry and enzyme translocation represent critical goals for mechanistic understanding of transcription. An essential aspect of reaching this goal is reliable capture of the Pol II active site with bound matched substrates in interaction with the TL. This has been difficult for the entire field. Here we demonstrate new crystallization conditions allowing capture of the TL in the closed state with matched substrates and couple this advance with next generation synchrotron technology. This technology has revealed aspects of the Pol II active site that have not previously been observed.

During crystallography experiments, the metal centers of metalloenzymes, such as the Mg ions in Pol II, can be susceptible to X-ray induced photo-reduction, which limits our ability to visualize important structural details and poses risks for the misinterpretation of structural results. Our results demonstrate the power of femtosecond diffraction methods at the XFEL to reveal functionally accurate structures of radiation sensitive metal centers, which eluded characterization at the synchrotron. Further, by using an efficient fixed-target approach for SFX data collection, we obtained these structures using only a few hundred crystals.

Our studies finally reveal the putative positions of Mg^2+^ A and B in the Pol II active site while additionally revealing a third Mg^2+^ ion. A third metal ion has been increasingly observed in many polymerases and nucleases previously proposed to function using two-metal ion catalysis^61,63,72,75-78^. Commonly, new technology has been required to observe these metals, such as time-resolved structural studies that can detect transient association. Here, use of XFEL was required to reveal both Mg^2+^ B and C in Pol II. What is function of this third metal and is it conserved in polymerases/nucleases? Most discussion has focused on DNA pol 17 and β, with proposals deriving from kinetics and structural observation that the function is catalytic in pol η^62,79^, while potentially product-stabilizing in pol β^63^. There is active discussion in the field about possible roles^78^, including some skeptics^80,81^. However, recent data strongly suggest arrival of third Mg2+ in pol η active site prior to chemistry in the case of a misincorporation event^79^.

What is the role of the third Mg^2+^ in Pol II? Deprotonation of ribose 3′-OH is a critical step for nucleophilic attack of the incoming NTP’s alpha phosphate. QM/MM simulations have shown that 3′-OH deprotonation is only energetically feasible if it is carried out by an active site water acting as a base^82^. It is especially difficult to pinpoint a potential acceptor for this proton given that it could be either an essential component for catalysis or a water^83^. Moreover, these studies showed that the presence of a third Mg^2+^ in the active site significantly lowered the activation energy and contributed to stabilization of the leaving pyrophosphate. Intriguingly, in other simulations, arrival of the third Mg^2+^ occurred prior to the nucleophilic attack of the α phosphate^84^, consistent with time-resolved crystallography of a pol η misincorporation event^79^. The latter could explain the presence of site C metal in both WT and 834P XFEL 3′ terminated structures that encompass an abortive reactant state with the active site residues and the three metals poised for substrate addition. Thus, the position of site C metal within 3 Å of the α phosphate in these structures could possibly depict its possible role in a path towards catalysis.

Mg^2+^ C is positioned adjacent to Rpb2^E529^ and Rpb2^Y769^. Alteration to Pol II by Rpb1^T834P^ shifts position of Mg^2+^ C towards the alpha phosphate andRpb2^Y769^ and away from Rpb2^E529^. Mutations to Rpb2^E529^ can increase (E529A/D) or decrease (E529Q) Pol II activity ^70^; similarly, mutation to Rpb2^Y769^ can alter Pol II activity (Y769F increases activity, Y769A predicted to decrease activity based on in vivo phenoypes^68^).

A large number of Pol II substitutions alter Pol II activity, including those that increase catalysis and perturb fidelity^23,25-27,43,67-70^. The majority, if not all, of increased activity Pol II mutants are within or in domains that surround the TL, supporting the idea that the TL conformation is finely balanced, and this balance can be altered by interactions with adjacent domains. These results raise the possibility that the TL is the endpoint for allosteric pathways that alter Pol II catalysis or translocation. Here, we have examined how the BH might contribute to Pol II activity and active site structure by analysis of Rpb1^T834P^. Mutational perturbation to the BH had consequences on structure and dynamics of the Pol II active site consistent with coupling of potential BH motion in WT to TL conformations and beyond. TL conformational changes effected by BH changes could both be direct through local contact changes such as disruption of the H-bond between T1077 to T834 in the proline mutant (**Fig. 5A**) or indirect through widening of the Pol II cleft. Deletion of Rpb9 alters Pol II properties and this has been interpreted through Rpb9 organization of adjacent Rpb1 domains that can stabilize the TL in the out/off conformation^67^. Cleft widening may function as a release or weakening of intra-Rpb1 interactions that stabilize TL in the out conformation. MD simulations indicate that in general the observed Mg^2+^ C is stably associated with the active sites of both WT and T834P Pol II but is closer to the alpha-phosphate oxygen (2.3 vs 2.9 A) and has altered coordination in the mutant (**Fig. 5B**). T834P Pol II shows greater BH-TL distances that appear to relate to increased flexibility in the T834P TL, consistent with coupling between the BH and the TL.

Key open questions in Pol II mechanism will be addressed by the incorporation of the time dimension in the catalytic cycle and use of systems that are able to observe translocation. Direct observation of states that promote or capture TL closing, and the relationship between TL-substrate interactions and chemistry will be required to reveal Pol II transcription at atomic resolution. Partially folded TL states have been observed and individual TL residues can be genetically distinguished, suggesting that the TL may function stepwise to promote catalysis. The order of these steps and when chemistry happens are unclear. Furthermore, translocation has not yet been structurally observed.

Time-resolved crystallography and cryo-EM are the approaches that likely will enable answering these types of questions. Cryo-EM will be especially valuable as examination of structure ensembles may identify translocation intermediates. Comparison of WT enzymes with mutants at the finely-detailed structural level, and for both matched NTP and mismatched substrates or with Mn^2+^ in place of Mg^2+^ will have the power to reveal plasticity of Pol II active site and give insight into evolution of RNA polymerase activity among eukaryotic and prokaryotic multisubunit RNAPs.

## Acknowledgements

This work was supported by NIH grants R35GM126948 to MF, R01GM097260 and R35GM144116 to CDK, and R01GM112686 to GC.

## Methods

### Protein Expression, Purification, and Complex Assembly

Pol II was purified as described^85^. In order to assemble a 3′-deoxy elongation complex (EC), we first incubated Pol II with 2.5 molar excess DNA/RNA scaffold in the presence of 5 mM MgCl_2_. A size exclusion chromatography (SEC) step was used to exchange buffer from the sample and remove the excess DNA/RNA scaffold from the sample. The eluted fractions were concentrated to 10 mg/ml for crystallization trials.

### Crystallization

Crystallization was performed using 10%-15% tacsimate and 100 mM HEPES pH 7.5. Crystal hits were generally obtained after 8-12 days. However, they were small (<50 µm), and diffracted to 5 Å resolution. Thus, to improve crystal morphology we applied a seeding protocol developed in the laboratory that allowed consistent diffraction to 3.3 Å as tested at beamline 12-2, SSRL ^86^This seeding protocol allowed crystal size optimization by modifying the protein: precipitant ratio. Crystals were cryoprotected in stages by increasing tacsimate in three steps from 10-15% to 60% in the span of three days; at the last step 10% glycerol was added to the buffer. Crystals were transferred to a 3 µL drop containing cryo-protectant solution (60% tacsimate and 10% glycerol, 100mM Hepes pH7.5), 2 mM ATP and 2mM Mg^2+^ (or Mn^2+^) and flash frozen in liquid nitrogen. Crystals for WT and T834P were mounted on custom grids developed in our laboratory to optimize data collection, with the strategy described in ^87^.

### Structure Refinement and Analysis

Diffraction experiments were performed using an 8 mm beam (40 fs duration) at 9.5 keV. Images were acquired using the Rayonix MX325 detector. Diffraction data was processed using the cctvx.xfel software package. The structures were solved by molecular replacement using MOLREP. Model building and refinement were performed using Coot and Phenix. The structures have been deposited in the Protein Data Bank.

### In vitro transcription

In vitro transcription for misincorporation was as described previously^25^ using template DNA CKO213, non-template DNA CKO435, with RNA9 preassembled into an elongation scaffold with the following exceptions. In vitro transcription buffer (20 mM Tris-HCl pH 8.0, 40 mM KCl, 5 mM MgCl_2_, 1 mM DTT) was supplemented to 10% glycerol and 0.25 mg/mL acetylated BSA (Ambion).

### MD Simulations

MD simulations for the WT and T834P mutant structures were performed. Missing loops with less than 8 residues were constructed using MODELLER version 9.15^88^. Histidine residue at 1085 was protonated based on the study by Huang et al^29^. The systems were solvated using cubic boxes with a 10 Å cutoff between the box edges and any atoms of the Pol II complexes. Box sizes were 158 Å for the WT system with the number of atoms of 391,755 and 160 Å for the Pol II T834P mutant with the number of atoms of 399,863. The systems were neutralized using Na^+^ atoms. Simulations were performed using CHARMM36m^89^ protein force field and CHARMM36 nucleic acid parameters^90,91^. TIP3P^92^ water model was used to solvate the systems. Modified charges were used for ATP and the terminal RNA residue as described in detail previously^93^. The systems were subjected to the energy minimization using 50 steps of steepest descent algorithm and 5000 steps of Adopted Basis Newton-Raphson algorithm. A total of 400 ps equilibration was performed by gradually increasing temperature from 5 to 298K and reducing the constraints on all the heavy atoms in the backbone and sidechains of proteins, P atoms of the nucleic acids and Mg atoms with a force constant from 10 kcal/mol/Å^2^ to 0 to relax the system completely. 100 ns of production runs with 5 replicates were run for each system that gives a total of 1 µs simulations. The simulations were performed using Langevin dynamics with a friction coefficient of 0.01 ps^-1^ under the constant temperature of 298 K. Periodic boundary conditions were used and particle mesh Ewald Algorithm was applied for the calculation of long-range electrostatic interactions. Lennard-Jones interactions were switched between 8 to 9 Å. The SHAKE algorithm was used to constraint bond lengths involving H atoms. Simulations were run using CHARMM version c43a1^91^ in conjunction with OPENMM^94^ using GPU machines. The Multiscale Modeling Tools for Structural Biology (MMTSB) tool set was used to carry out simulations and analysis^95^. Clustering of the structures stored in the trajectories was performed using k-clustering approach based on the Time-structure Independent Component Analysis (TICA) reduced dimension of pairwise residue distances for the TL residues. This analysis was performed using MSMBuilder version 3.8^96^. Mg^2+^ density analysis was performed on the reoriented trajectories in such a way that the active site of the systems that contains BH and TL residues and the nucleic acids including RNA, DNA and ATP were aligned to the crystal structures for every frame. The densities for Mg^2+^ were calculated on a three-dimensional grid with 1 Å resolution throughout the trajectories of five replicates for each system.

## Supplementary Figure Legends. See word document (uploaded in T834P)

**Supp. Table 1.** Data Collection and refinement statistics.

|  | WT LCLS (MFX) | T834P LCLS (XPP) | WT-Mncl2 (SSRL)<sup>a</sup> | WT-NoATP (SSRL)<sup>a</sup> |
| --- | --- | --- | --- | --- |
| **Data collection** |  |  |  |  |
| Number of Crystal | 320 | 220 | 1 | 1 |
| Total number of Images | 650 | 1085 |  |  |
| Indexed/Merged (%of total images) | 480 (73.8) | 883(81.4) |  |  |
| Space group | C222<sub>1</sub> | C222<sub>1</sub> | C222<sub>1</sub> | C222<sub>1</sub> |
| a, b, c (Å) | 217.20, 387.44, 276.96 | 220.17, 392.60, 280.92 | 221.19, 393.97, 283.3 | 219.09, 390.92, 281.65 |
| $\alpha$, $\beta$, $\gamma$ (°) | 90, 90, 90 | 90, 90, 90 | 90, 90, 90 | 90, 90, 90 |
| Resolution (Å) | 24.68-3.30 | 24.96-3.0 |  |  |
| R<sub>merge</sub> | 59.99 (74.0)<sup>b</sup> | 66.74(47.7) |  |  |
| I/σI | 3.93(0.36) | 4.91 (1.65) | | |
| Completeness (%) | 95.85 (67.53) | 99.2 (95.1) |  |  |
| Redundancy | 10.25 (2.18) | 8.82(4.25) |  |  |
| CC1/2 | 91.44(59.0) | 75.37 (20.11) |  |  |
| **Refinement** |  |  |  |  |
| Resolution (Å) | 19.97-3.34 | 18.00-3.05 | 49.25-3.40 | 35.08-3.51 |
| No. reflections | 165059 | 223774 | 150237 | 133630 |
| R<sub>work</sub>/R<sub>free</sub> | 25.51/26.13 | 28.3/30.7 | 22.01/24.87 | 21.86/24.56 |
| **No. atoms** |  |  |  |  |
| Protein | 30109 | 30236 | 30107 | 30252 |
| RNA/DNA | 496 | 477 | 477 | 458 |
| other | 48 | 48 | 14 | 8 |
| Water | 18 | 124 | 142 | 0 |
| **R.m.s deviations** |  |  |  |  |
| Bond lengths (Å) | 0.002 | 0.011 | 0.003 | 0.003 |
| Bond angles (°) | 0.593 | 1.554 | 0.670 | 0.712 |
| **PDB:ID** | 8U9R | 8U9X |  |  |

<sup>a</sup>One crystal was used for data collection and structure determination.
<sup>b</sup>Values in parentheses are for the highest resolution shell.

**Supplementary Figure 1.**
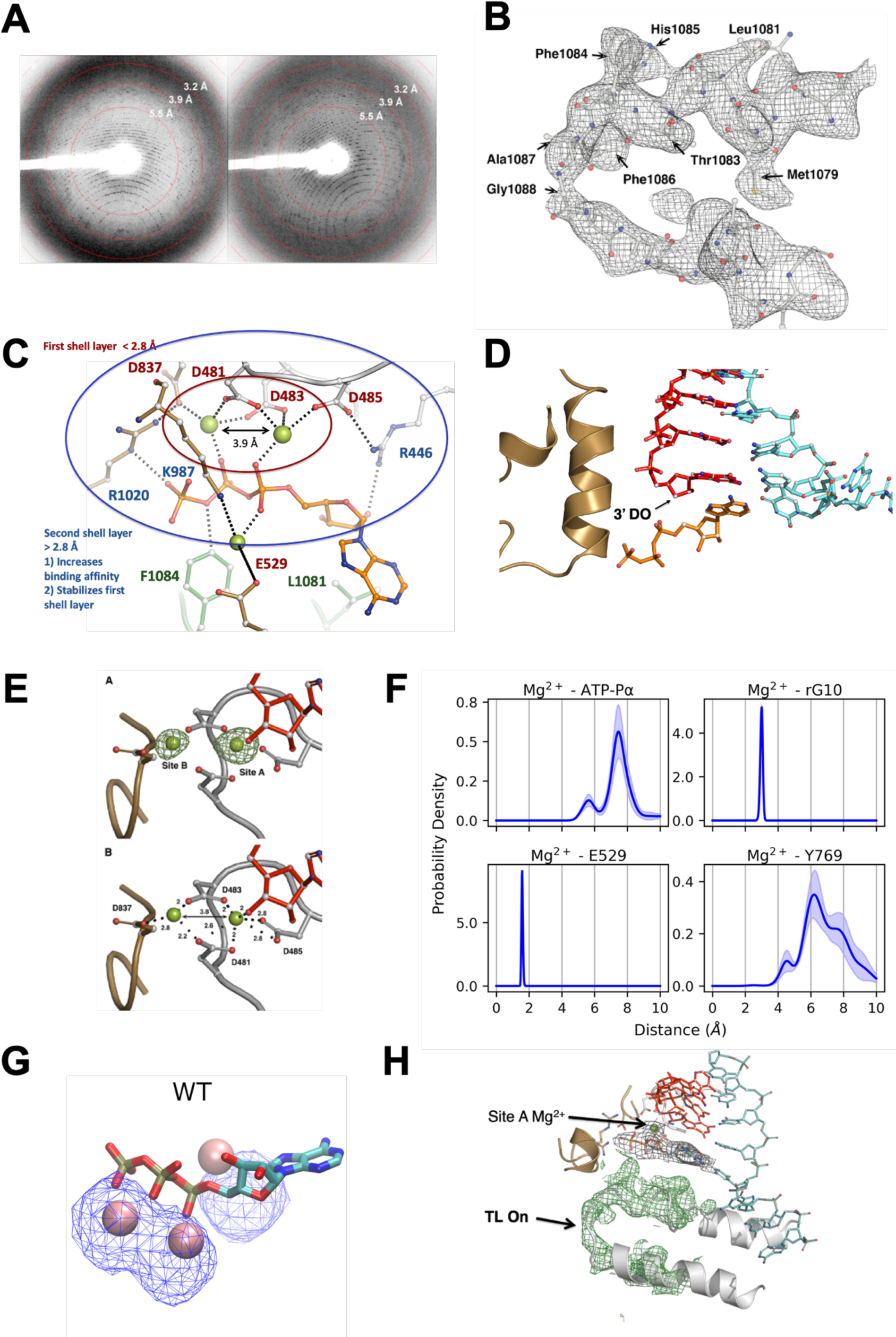
**A**. Two diffraction XFEL images. **B**. Fo-Fc (difference) map of TL contoured at 3 σ. **C**. Magnesium coordination shells in Pol II active site. **D**. Active site – A chain terminating nucleotide lacks the ribose 3′-OH. **E**. Top panel. Green mesh, electron density map countered at 10 α for metals A and B for the 250 mM MnCl_2_ soak. Bottom panel. Coordination of metal sites A and B, numbers correspond to distance in angstroms. **F**. Active site – MD simulations – Probability density distributions of distances of Mg^2+^ at the C site to the P_α_ of ATP, terminal RNA residue (rG10), Rbp2 E529 and Y769, the errors between five replicate simulations were provided as shaded areas. **G.** Distribution of Mg^2+^ ion calculated over MD simulation trajectories of WT Pol II, density was shown as isosurface at 1.0% occupancy. **H**. Green mesh, electron density map countered at 3 α for the closed state of the TL. Data was collected using conventional synchrotron radiation (to 3.3 Å) using crystals from the same batch to those collected using XFEL. Only site A metal was detected.

**Supplementary Figure 2.**
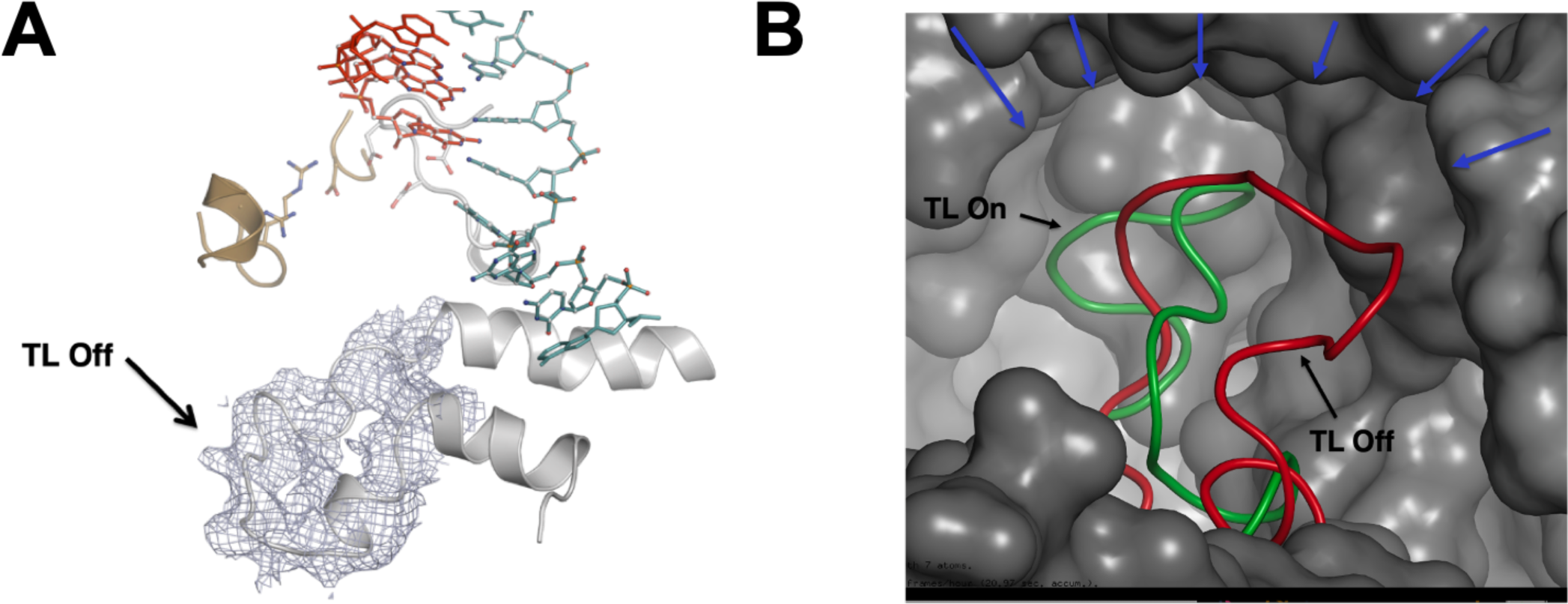
**A**. Grey mesh, electron density map countered at 2.5 α for the TL in open conformation. No substrate was soaked in the crystals. **B**. Physical path formed by Rpb1 and Rpb2 residues restricting TL motion during open and closed states.

**Supplementary Figure 3.**
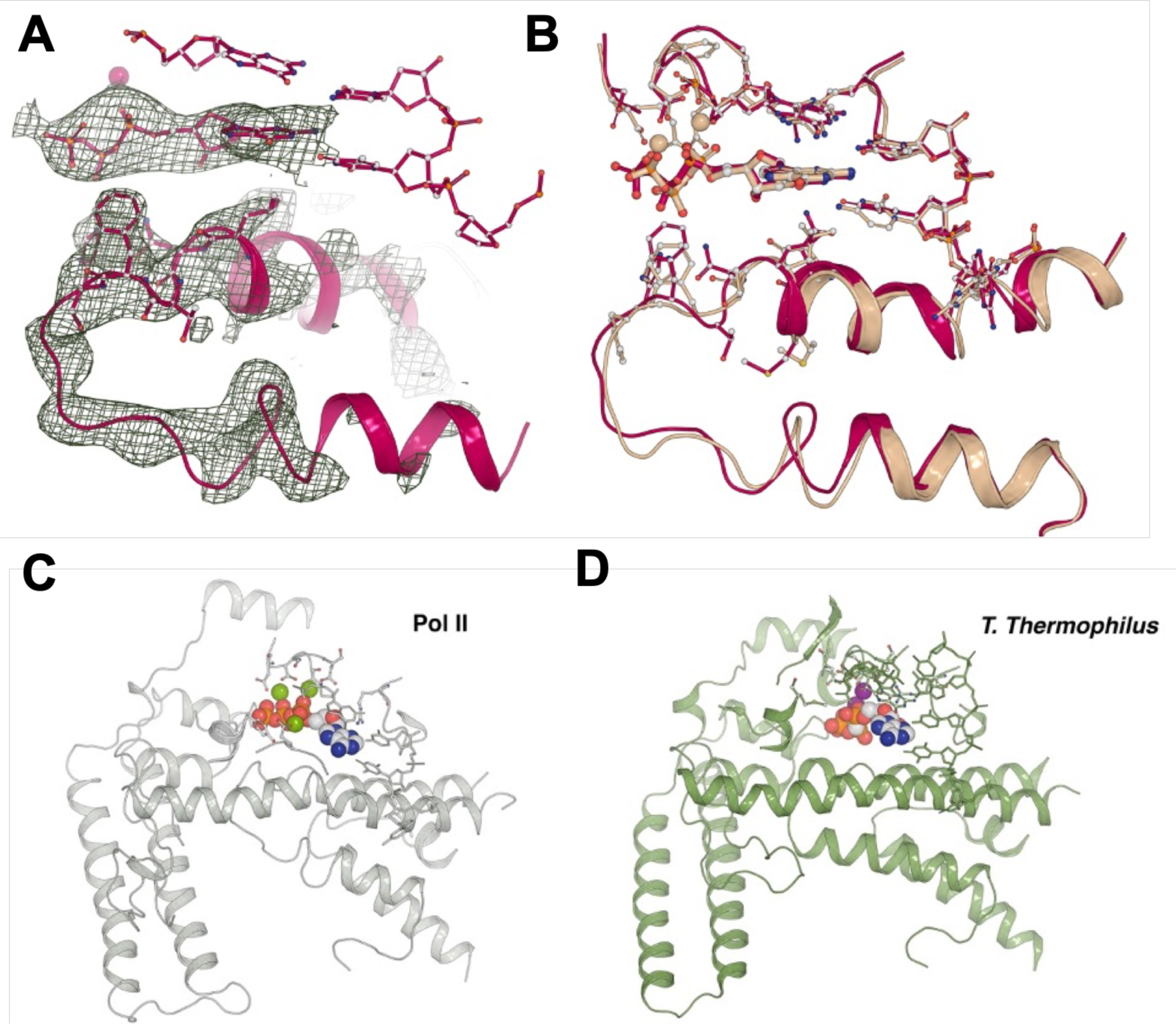
**A** Grey mesh, Fo-Fc electron density map countered at 2.5 σ for the TL in the re-refined TL in closed conformation. **B**. Overlay of the 2E2H structure (light pink) and the re-refined 2E2H (dark pink), the clash score after refinement improved from 156 to 17. **C and D.** Active site comparison between Pol II and *TTh* illustrating analogous architectures.

**Supplementary Figure 4.**
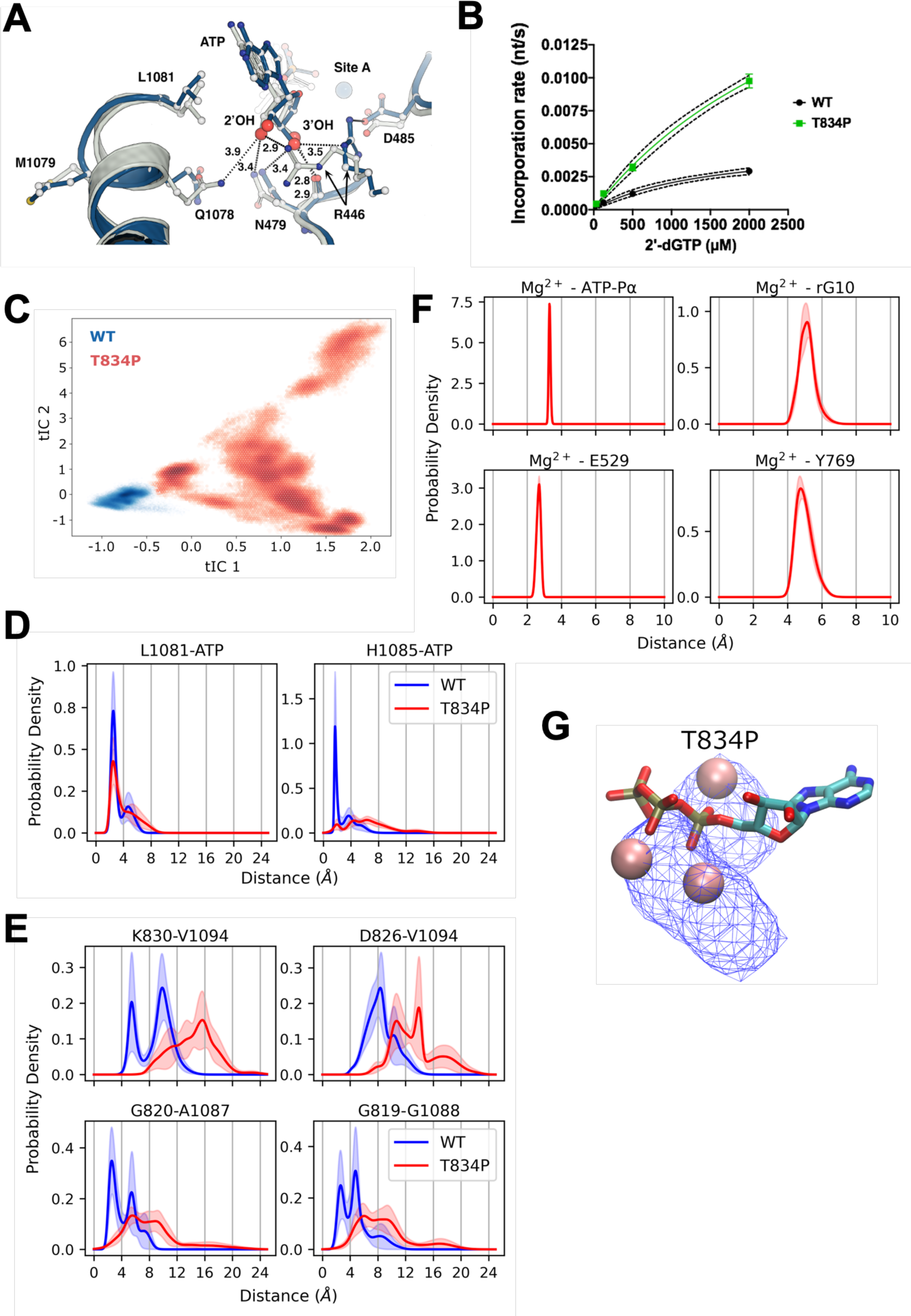
**A** Overlay of WT (silver) vs T834P (blue) structures to illustrate residue H-bond formation with 2′-OH and 3′-OH (large red spheres). H-bond contacts with the 2′-OH for Q1078 R446 and N479 are lost in the mutant. **B.** Increased misincorporation rate for T834P Pol II for addition of a single 2′-dGTP on a template specifying addition of ATP in vitro. **C.** Time-structure independent component analysis (TICA) based on pairwise minimum distances of TL residues for WT and T834P simulations. **D-F.** Probability density distributions of distances between TL residues (L1081 and H1085) and ATP for WT and T834P simulations (D), and between TL and BH residues for WT and T834P simulations (E) or between Metal C and active site features for T834P simulations (F, compare with Sup. Figure 1F). All the distances were calculated for five replicates and errors were provided as the shaded area. **G.** Distribution of Mg^2+^ ion calculated over MD simulation trajectories of T834P, density was shown as isosurface at 1.0% occupancy.

## Notes

### Competing Interest Statement

The authors have declared no competing interest.

